# Base-pairing requirements for small RNA-mediated gene silencing of recessive self-incompatibility alleles in *Arabidopsis halleri*

**DOI:** 10.1101/370239

**Authors:** N. Burghgraeve, S. Simon, S. Barral, I. Fobis-Loisy, A-C Holl, C. Poniztki, E. Schmitt, X. Vekemans, V. Castric

## Abstract

Small non-coding RNAs are central regulators of genome activity and stability. Their regulatory function typically involves sequence similarity with their target sites, but understanding the criteria by which they specifically recognize and regulate their targets across the genome remains a major challenge in the field, especially in the face of the diversity of silencing pathways involved. The dominance hierarchy among self-incompatibility alleles in Brassicaceae is controlled by interactions between a highly diversified set of small non-coding RNAs produced by dominant S-alleles and their corresponding target sites on recessive S-alleles. By controlled crosses, we created numerous heterozygous combinations of S-alleles in *Arabidopsis halleri* and developed an RT-qPCR assay to compare allele-specific transcript levels for the pollen determinant of self-incompatibility (*SCR*). This provides the unique opportunity to evaluate the precise base-pairing requirements for effective transcriptional regulation of this target gene. We found strong transcriptional silencing of recessive *SCR* alleles in all heterozygote combinations examined. A simple threshold model of base-pairing for the sRNA-target interaction captures most of the variation in *SCR* transcript levels. For a subset of S-alleles, we also measured allele-specific transcript levels of the determinant of pistil specificity (*SRK*) and found sharply distinct expression dynamics throughout flower development between *SCR* and *SRK*. In contrast to *SCR*, both *SRK* alleles were expressed at similar levels in the heterozygote genotypes examined, suggesting no transcriptional control of dominance for this gene. We discuss the implications for the evolutionary processes associated with the origin and maintenance of the dominance hierarchy among self-incompatibility alleles.

## Introduction

Small non-coding RNAs are short RNA molecules (20-25nt) with a range of regulatory functions (Vazquez *et al*., 2010; Aalto & Pasquinelli, 2012). The best-known members of this class of molecules are microRNAs, which are typically involved in post-transcriptional gene silencing and regulate the activity of their target gene in *trans* by either mRNA cleavage (quickly followed by degradation) or by blocking translation (Li *et al*., 2014). In some cases, the action of microRNAs leads to the production of secondary phased short interfering RNAs (pha-siRNAs) by their target coding or non-coding sequence, which in turn can regulate other downstream targets (Fei *et al*., 2013). Another major set of small RNAs is heterochromatic short interfering RNAs (hc-siRNAs) which are mediating transcriptional silencing of repeat sequences in the genome through epigenetic modification by the RNA-dependent DNA methylation pathway (RdDM, Matzke *et al*., 2009).

Both microRNAs and siRNAs guide their effector molecules (members of the ARGONAUTE gene family: AGO1 and AGO4, respectively) to their target sites by sequence similarity through base-pairing. For plant microRNAs, sequence similarity with the target sequence is typically very high and appears to be a shared feature of all functionally verified interactions (Wang *et al*., 2015). High base-pairing complementarity, however, is not the sole determinant of target specificity, and the position of the mismatches along the microRNA:target duplex is also important. Indeed, expression assays showed that while individual mismatches typically have limited functional consequences, they can also entirely inactivate the interaction when they hit specific positions such as, for example, the 10^th^ and 11^th^ nucleotide, corresponding to the site of cleavage (Jones-Rhoades *et al*., 2006). Furthermore, the position of mismatches along the microRNA:target duplex also seems to be crucial, with a greater tolerance in the 3’ than the 5’ region of the microRNA (up to four mismatches generally have limited functional consequences in the 3’ region, while only two mismatches in the 5’ region seem sufficient to abolish the target recognition capability; Liu *et al*., 2014, Mallory *et al*., 2004; Parizotto *et al*., 2004; Schwab *et al*., 2005). These observations have led to the formulation of general “rules” for microRNA targeting (Axtell & Meyers, 2018), but at the same time they also revealed a large number of exceptions. As a result, *in silico* prediction of microRNA target sites currently remains a difficult challenge (Ding *et al*., 2012; Axtell & Meyers, 2018). For other types of small RNAs (pha-siRNAs and hc-siRNAs), even less is known about the base-pairing requirements for targeting, mostly because of the absence of experimentally confirmed examples of discrete, single siRNA target sites either in *cis* or in *trans* (Wang *et al*., 2015).

In this context, the recent discovery by Tarutani *et al*. (2010), Durand *et al*. (2014) and Yasuda *et al*., (2016) of a highly diversified set of small non-coding RNAs at the gene cluster controlling self-incompatibility (SI) in Brassicaceae, provides an experimentally tractable model to evaluate the base-pairing requirements for silencing by a set of sRNAs that are regulating expression of a single gene. Sporophytic SI is a genetic system that evolved in several hermaphroditic plant lineages to enforce outcrossing by preventing self-fertilization, hence avoiding inbreeding depression (De Nettancourt, 2001). In the Brassicaceae family, SI is controlled by a single genomic region called the “S-locus”, which contains two tightly linked genes, namely *SCR* and *SRK*, that encode the pollen S-locus cysteine-rich and the stigma S-locus receptor kinase recognition proteins, respectively. This system involves a polymorphism in which multiple deeply diverged allelic lines are maintained, and accordingly a large number of S-alleles is typically found in natural populations of self-incompatible species (Castric & Vekemans, 2004). With such a large allelic diversity and the very process of self-rejection, most individual plants are heterozygotes at the S-locus. Yet in most cases, only one of the two S-alleles in a heterozygous genotype is expressed at the phenotypic level in either pollen or pistil, as can be revealed by controlled pollination assays on pollen or pistil tester lines (Llaurens *et al*., 2008; Durand *et al*., 2014). Which of the two alleles is expressed is determined by their relative position along a dominance hierarchy, whose molecular basis for the pollen phenotype has been initially studied in the genus Brassica. In this genus, dominance is controlled at the transcriptional level in pollen (Schopfer 1999, Kakizaki *et al*. 2003). Transcriptional silencing of recessive alleles by dominant alleles is caused by 24nt-long trans-acting small RNAs produced by dominant S-alleles and capable of targeting a DNA sequence in the promoter sequence of the *SCR* gene of recessive S-alleles, provoking DNA methylation (Shiba *et al*. 2006). Details of how these sRNAs achieve their silencing function remain incompletely understood (Finnegan *et al*., 2011), but it is clear that their biogenesis is similar to that of microRNAs (*i.e.*, they are produced by a short hairpin structure), while their mode of action is rather reminiscent of that of siRNAs (*i.e.*, the transcriptional gene silencing functions through recruitment of the methylation machinery). Strikingly, the full dominance hierarchy in the Brassica genus seems to be controlled by just two small RNAs called *Smi* and *Smi2* (Tarutani *et al*., 2010, Yasuda *et al*. 2016). *Smi* and *Smi2* target distinct DNA sequences, but both are located in the promoter region of *SCR*, and both seem to involve DNA methylation and 24-nt active RNA molecules.

The dominance hierarchy in Brassica is, however, peculiar in that only two ancestral allelic lineages segregate in that genus (the class I and class II alleles referred to above, see *e.g.* Leducq *et al*., 2014), whereas self-incompatible species in Brassicaceae typically retain dozens of highly divergent ancestral allelic lineages (Castric & Vekemans, 2004). A recent study showed that in *Arabidopsis halleri*, a Brassicaceae species with multiple allelic lineages at the S-locus, the dominance hierarchy among S-alleles in pollen is controlled by not just two but as many as eight different sRNA precursor families and their target sites, whose interactions collectively determine the position of the alleles along the hierarchy (Durand *et al*., 2014). In that genus, much less is known about the mechanisms by which the predicted sRNA-target interactions translate into the dominance phenotypes. First, the expression dynamics of the *SCR* gene across flower development stages is poorly known. Indeed, Kusaba *et al*. (2002) measured expression of *SCR* alleles in *A. lyrata*, but focused on only two S-alleles (*SCRa* and *SCRb*, also known as *AlSCR13* and *AlSCR20*, respectively, in Mable *et al*. 2003) and showed striking differences in their expression dynamics in anthers. Hence, the developmental stage at which the transcriptional control of dominance in pollen should be tested is not precisely known. Second, while they did confirm monoallelic expression, consistent with the observed dominance relationship between the two alleles (*SCR*b > *SCR*a, Kusaba *et al*. 2002), the fact that only a single heterozygote combination was measured among the myriad possible combinations given the large number of S-alleles segregating in that species (at least 43 S-alleles: Genete *et al*., 2020) prevents generalization at this step. Hence, a proper experimental validation of the transcriptional control of dominance among S-alleles in the Arabidopsis genus is still lacking. Third, Durand *et al*., (2014) observed rare sRNA-target interaction predictions that did not agree with the observed dominance phenotype. In particular, they identified pairs of S-alleles where no sRNA observed as being produced by the dominant allele was predicted to target the *SCR* gene of the recessive one, while the dominance phenotype had been well established phenotypically by controlled crosses (*e.g.* Ah04>Ah03) suggesting the possibility that mechanisms other than transcriptional control may be acting.

Conversely, in other rare cases, sRNAs produced by a recessive S-allele were predicted to target the *SCR* gene of a more dominant allele, suggesting exceptions to the set of base-pairing rules used to predict target sites. Fourth, the target sites for the two sRNAs in Brassica were both located in the promoter sequence (Tarutani *et al*., 2010, Yasuda *et al*. 2016), and can thus reasonably be expected to prevent transcriptional initiation through local modification of the chromatin structure associated with DNA methylation. Many of the predicted sRNA target sites in *A. halleri*, however, are rather mapped to the *SCR* intron or the intron-exon boundary (beside some in the promoter as well, Durand *et al*. 2014), which suggests that distinct silencing pathways might be acting (Cuerda-Gil & Slotkin, 2016). It thus remains to be determined whether transcriptional control is also valid when the targets are at other locations along the *SCR* gene structure. Finally, the dominance hierarchy at the female determinant *SRK* differs from that at *SCR*, co-dominance being more frequent than on the pollen side both in Brassica (Hatakeyama *et al*., 2001) and in *A. halleri* (Llaurens *et al*., 2008). Limited transcriptional analysis in Brassica and Arabidopsis suggests that dominance in pistils is not associated with *SRK* expression differences, but again the number of allelic pairs tested has remained limited (Suzuki *et al*. 1999; Kusaba *et al*. 2002).

Here, we take advantage of the fact that dominance interactions in Arabidopsis SI are controlled in pollen by a diversity of sRNAs and the diversity of their target sites to determine the base-pairing requirements for successful small-RNA mediated transcriptional silencing of recessive *SCR* alleles. We first used controlled crosses to obtain a large collection of *A. halleri* plants in which S-alleles were placed in various homozygous and heterozygote combinations for which pairwise dominance interactions had been determined. We then developed and validated a qPCR protocol for allele-specific expression of a set of nine *SCR* and five *SRK* alleles in *A. halleri*. This enabled us to analyse the expression dynamics across four flower developmental stages of each of these alleles and test the transcriptional control of dominance for both genes in many heterozygote combinations. We quantified the strength of silencing of recessive *SCR* alleles and propose a quantitative threshold model for how sequence identity between the small non-coding RNAs and their target sites results in silencing. We discuss the implications of this model on the evolutionary processes associated with the origin and maintenance of the S-locus dominance hierarchy in Brassicaceae.

## Material & Methods

### Plant material

We used controlled crosses to create a collection of 88 *A. halleri* plants containing nine different S-alleles (S1, S2, S3, S4, S10, S12, S13, S20, and S29) in a total of 37 of all 45 possible homozygous and heterozygous combinations. Some *S*-locus genotypes were obtained independently by different controlled crosses and were considered below as “biological replicates” (different genetic backgrounds, on average *n*= 2.05 biological replicates per S-locus genotype, Table S1 & S2). Three plants were cloned by cuttings and considered as “clone replicates” (identical genetic background, Table S1) that we used to evaluate the expression variance associated with different genetic backgrounds.

Each plant was genotyped at the S-locus using the PCR-based protocol described in Llaurens *et al*. (2008). Pairwise dominance interactions between S-alleles of the heterozygote combinations were either taken from Llaurens *et al*. (2008); Durand *et al*. (2014); Leducq *et al*. (2014) or were newly determined by controlled pollination assays following the protocol of Durand *et al*., (2014). In a few instances, relative dominance status of the two alleles had not been resolved phenotypically and were inferred from the phylogeny of *SRK* alleles, which is largely consistent with the dominance hierarchy (Durand *et al*. 2014). The pairwise dominance interactions between these alleles as determined by pollen and pistil compatibility phenotypes of heterozygote plants are reported in Table S3.

### RNA extraction and reverse transcription

On each plant, we collected flower buds at four developmental stages: 1) five highly immature inflorescence extremities (more than 2.5 days before opening, buds below 0.5mm, stages 1-10 in *A. thaliana* according to Smyth *et al*., 1990); 2) ten immature buds (2.5 days before opening, between 0.5 and 1mm, approximately stage 11); 3) ten mature buds (one day before opening, longer than 1mm, approximately stage 12); and 4) ten open flowers (approximately stages 13-15). These stages were characterized by establishing the size distribution within each stage and measuring the time to flower opening based on ten buds. Samples collected were flash-frozen in liquid nitrogen, then stored at −80°C before RNA extraction. Tissues were finely ground with a FastPrep-24 5G Benchtop Homogenizer (MP Biomedicals, Model #6004-500) equipped with Coolprep 24 × 2mL adapter (6002-528) and FastPrep Lysis Beads & Matrix tube D. Total RNAs were extracted with the Arcturus “Picopure RNA isolation” kit from Life Science (PN: KIT0204) according to the manufacturer’s protocol, including a step of incubation with DNAse to remove gDNA contamination. We normalized samples by using 1 mg of total RNA to perform reverse-transcription (RT) using the RevertAid Fermentas enzyme following the manufacturer’s instructions.

### Primer design

A major challenge to study expression of multiple S-alleles is the very high levels of nucleotide sequence divergence among them, precluding the possibility of designing qPCR primers that would amplify all alleles of the allelic series (both for *SRK* and *SCR*). Hence, we rather designed qPCR primers specifically targeted towards each of the *SCR* and *SRK* alleles, and for each heterozygote genotype we independently measured expression of both alleles of each gene. Primers were designed based on genomic sequences from BAC clones (Goubet *et al*. 2012; Durand *et al*. 2014; Novikova *et al*. 2017), with a length of ∼20 nucleotides, a GC content around 50% and a target amplicon size around 150nt (Figure S1). For *SCR*, we focused on a set of 9 S-alleles. Whenever possible, we placed primers on either side of the *SCR* intron to identify and discard amplification from residual gDNA. However, because the coding sequence of the *SCR* gene is short, the number of possible primers was limited and this was not always possible. In two cases (*SCR01* and *SCR20*), both primers were thus located within the same exon. For *SRK* alleles, the primers were also designed on either side of the first intron to avoid genomic contamination (Figure S2).

Because no differences in transcript levels were previously observed between dominant and recessive *SRK* alleles (Suzuki *et al*. 1999; Kusaba *et al*. 2002), and given the effort required to optimize new qPCR primers, we decided to place more effort on *SCR* and focused on a more limited number of *SRK* alleles (*n*=5). To obtain relative expression levels across samples, we used *actin 8* (At1g49240) as a housekeeping gene for standardization after we verified that the *A. thaliana* and *A. halleri* sequences are identical at the primer positions (An *et al*. 1996). Primer sequences are reported in Table S4.

### Quantitative real-time PCR

On each cDNA sample, at least three qPCR reactions (referred to below as “technical” replicates) were performed for *actin 8* and for each of the S-alleles contained in the genotype (one S-allele for homozygotes, two S-alleles for heterozygotes). The runs were made on a LightCycler480 (Roche) with iTaq Universal SYBR Green Supermix (Bio-rad, ref 172-5121). Amplified cDNA was quantified by the number of cycles at which the fluorescence signal was greater than a defined threshold during the logarithmic phase of amplification using the LightCycler 480 software release 1.5.0 SP3. The relative transcript levels are shown after normalisation with actin amplification through the comparative 2^−Δ*Ct*^ method (Livak & Schmittgen, 2001). The *Ct*_SCR_ and *Ct*_SRK_ values of each technical replicate were normalized relative to the average *Ct*_actin_ measure across the three replicates.

### Validation of qPCR primers at the dilution limits

Given the very large nucleotide divergence between alleles of either *SCR* or *SRK*, cross-amplification is unlikely. However, to formally exclude that possibility, we first performed cross-amplification experiments by using each pair of *SCR* primers on a set of cDNA samples that did not contain that target *SCR* allele but instead contained two other *SCR* alleles in various heterozygous genotypic combinations (*n*=7 on average). In order to evaluate our ability to measure expression of *SCR* alleles in biological situations where they are expected to be transcriptionally silenced, we then used a series of limit dilutions to explore the loss of linearity of the relationship between *Ct* and the dilution factor (six to eight replicates per dilution level). Then we examined the shape of the melting curves to determine whether our measures at this limit dilution reflected proper PCR amplification or the formation of primer dimers. Finally, we used water in place of cDNA to evaluate the formation of primer dimers in complete absence of the target template DNA.

### Expression dynamics and the effect of dominance

We used generalized linear mixed models (lme4 package in *R*; Bates *et al*., 2014) to decompose *Ct* values normalized by the *actin 8* control (as the dependent variable) into the effects of five explanatory variables. Two of them were treated as fixed effects: developmental stage (4 categories) and relative dominance of the allele studied in the genotype (3 categories: recessive, dominant, homozygous). Because expression of the different *SCR* (and *SRK*) alleles was quantified by different primer pairs with inevitably different amplification efficiencies, *Ct* values cannot be directly compared across alleles and accordingly we included the identity of *SCR* or *SRK* alleles as random effects. Biological and clone replicates were also treated as random effects, with clones nested within biological replicates (Table S5). We visually examined normality of the residuals of the model under different distributions of 2^−Δ*Ct*^, including Gaussian, Gamma and Gaussian with logarithmic transformations. We tested whether the different *S*-alleles have different expression profiles across developmental stages, as suggested by Kusaba *et al*. (2002) for *SCR* in *A. lyrata*, by using ANOVA to compare nested models in which a random effect for the interaction between the “allele measured” and “stage” effects was either absent (model 1) or introduced (model 2, Table S5b) in addition to the fixed effect of stage. The existence of this interaction was tested for *SCR* and *SRK* separately.

### Target features and silencing effect

Expression of *SCR* in heterozygote genotypes in *A. halleri* is controlled by a small RNA-based regulatory machinery (Durand *et al*. 2014). We then sought to determine how *SCR* transcript levels were affected by specific features of the small RNA-target interactions between *S*-alleles. We retrieved sRNA sequencing data from individuals carrying eight of the nine *S*-alleles considered (S01, S03, S04, S10, S12, S13 and Ah20 from Durand *et al*. (2014) and S02 from Novikova *et al*. (2017)). No sRNA sequencing data were available for the last *S*-allele (S29). We used these sRNA sequencing data to determine the complete set of sRNA molecules uniquely produced by the annotated sRNA precursors of each of these eight *S*-alleles. To do that, we mapped the sRNA reads to the sRNA precursor sequences carried by the respective S-alleles after excluding those that mapped to other locations in the closely related *A. lyrata* genome (Durand *et al*. 2014). For each sRNA produced by a given S-allele, we then predicted putative target sites on the *SCR* gene of all other S-alleles including 2kb of genomic nucleotide sequence both upstream and downstream of *SCR* using a dedicated alignment algorithm and scoring matrix, as described in Durand *et al*. (2014). Briefly, alignment quality was assessed by a scoring system based on the addition of positive or negative values for matching nucleotides (+1), mismatches and gaps (−1), taking into account the non-canonical G:U interaction (−0.5). For each pair of alleles considered, only the sRNA/target combination with the highest score was selected for further analysis (Table S6). The analysis was performed regardless of the dominance relationship (i.e. we predicted putative target sites of sRNAs produced by dominant S-alleles onto recessive S-alleles, and reciprocally from recessive S-alleles onto dominant S-alleles). Because the mechanisms by which silencing is achieved remain unclear at this stage, we did not filter these sRNA further in terms of length or identity of the 5’ nucleotide, in line with Durand *et al*. (2014). In the cases where the target with the highest score was due to a sRNA with non-canonical size (anything but 21 or 24nt), we also reported the best target score among the set of 21 and 24nt sRNA molecules produced by the same S-allele (Table S6). We used Akaike Information Criteria (AIC) to compare how well different base-pairing scores for target site identification predicted the level of *SCR* expression (and hence the silencing phenomenon), varying the threshold from 14 to 22. Lower values of AIC are associated with a best fit of the model. We then added a new fixed effect in our basal model to test whether targets at different positions along the *SCR* gene (5 categories: 5’ portion, exons, intron, overlapping the exon-intron boundary or 3’ portion of the gene) are associated with different strengths of silencing. For this analysis, we included only targets above the threshold identified (score >= 18).

Effective silencing of recessive *SCR* alleles in *Brassica rapa* is dependent upon combinations of individual sequence mismatches between the *Smi* & *Smi2* small RNAs and their target sites in the class II alleles (Yasuda *et al*., 2016), but interaction in this study relied on raw counts of nucleotide mismatches and were thus not directly comparable to our results. To determine whether the base-pair requirements for silencing are similar, we thus reanalysed these interactions using our scoring system to compare the small RNA-target alignment scores between Brassica and Arabidopsis (Tarutani *et al*., 2010, Yasuda *et al*., 2016).

Finally, we used the phylogeny in Durand *et al*. (2014) to classify sRNA/target interactions into “recent” (mir867 and mirS4) and “ancient” (mirS1, mirS2 mirS3, mirS5, mir1887 and mir4239). Based on this classification, we used a linear regression to compare the alignment score for recent and ancient sRNAs and tested the hypothesis that interactions with base-pairing scores above the threshold at which silencing was complete correspond to recently emerged interactions that have not yet accumulated mismatches.

## Results

### Validation of the qPCR protocol and the allele-specific primers

The specificity test confirmed the absence of cross-amplification between alleles, as the *Ct* measures for water control and cross amplification were comparably high (around *Ct*=34) and both were higher than the positive controls (median *Ct*=22, Figure S3). Overall, serial dilutions of the template cDNA confirmed linearity of the *Ct* measure within the range of values observed for a given allele across the different conditions examined (Figure S4a). Because we study a silencing phenomenon, we then explored how signal was lost at the dilution limits. As expected, linearity started to be lost at very low cDNA concentrations (in particular for alleles *SCR01*, *SCR02*, *SCR04*, *SCR13* and *SCR20*, Figure S4a), and examination of melting curves under these conditions indicated the formation of primer dimers rather than the expected transcripts. Hence, we note that comparing levels of expression for a given allele between different recessive contexts (*e.g.* when silenced by different sRNAs) should be challenging, especially for the above-mentioned alleles. Linearity was good for most *SRK* alleles (Figure S4b) except for *SRK12* (data not shown), so this allele was excluded from further analyses.

*SCR* and *SRK* expression dynamics across flower development stages In total, we performed 344 RNA extractions and RT-PCR from the 37 different S-locus genotypes sampled at four developmental stages. For *SCR*, we measured 1,838 *Ct_SCR_*/*Ct*_actin_ expression ratios (*i.e.* an average of 26.9 expression measures per S-allele in each diploid genotype, Table S1). For *SRK*, we measured 480 *Ct_SRK_*/*Ct*_actin_ ratios (*i.e.* an average of 11.1 expression measures per S-allele in each diploid genotype, Table S2). Distribution of the residuals of the generalized mixed linear model was closest to normality after log-transformation of the ratios (Figure S6). As expected, measured expression levels were more highly repeatable across clones than across biological replicates for a given S-locus genotype (deviance estimates of 0.40, 1.08, respectively, Table S5a). The deviance associated with the allele’s expression dynamic was higher (deviance = 4.56), although we note that the technical error was also important (deviance = 6.08, Table S5a). We first examined the expression dynamics of the different *SCR* alleles. Because recessive *SCR* alleles were consistently silenced (see below), we isolated the effect of developmental stages by focusing only on genotypes in which each focal allele was known to be dominant at the phenotypic level (Figure 1a). Overall, we observed a strong pattern of variation among stages (F-value: 10.76, *p*-value: 5.7e-5, Table S5c) with high expression of *SCR* in buds at early developmental stages (<0.5 to 1mm), and low expression in late buds right before opening and in open flowers. This pattern is consistent with degeneration in these stages of the anther tapetum, the cellular layer where *SCR* is expected to be expressed. The expression dynamics of *SRK* was sharply different from that of *SCR*, with monotonously increasing expression in the course of flower development, with lowest expression in immature buds (<0.5mm) and highest expression in open flowers (Figure 1b, F-value: 4.411, *p*-value: 0.007, Table S5h). We found evidence that the expression dynamics varied across *S*-alleles, not only for *SCR* (Chi^2^: 308.19, *p*-value < 2.2e-16, Table S5b) in line with Kusaba *et al*., (2002), but also for *SRK* (Chi²: 6.9103, *p*-value 0.00857, Table S5g).

**Figure 1:**
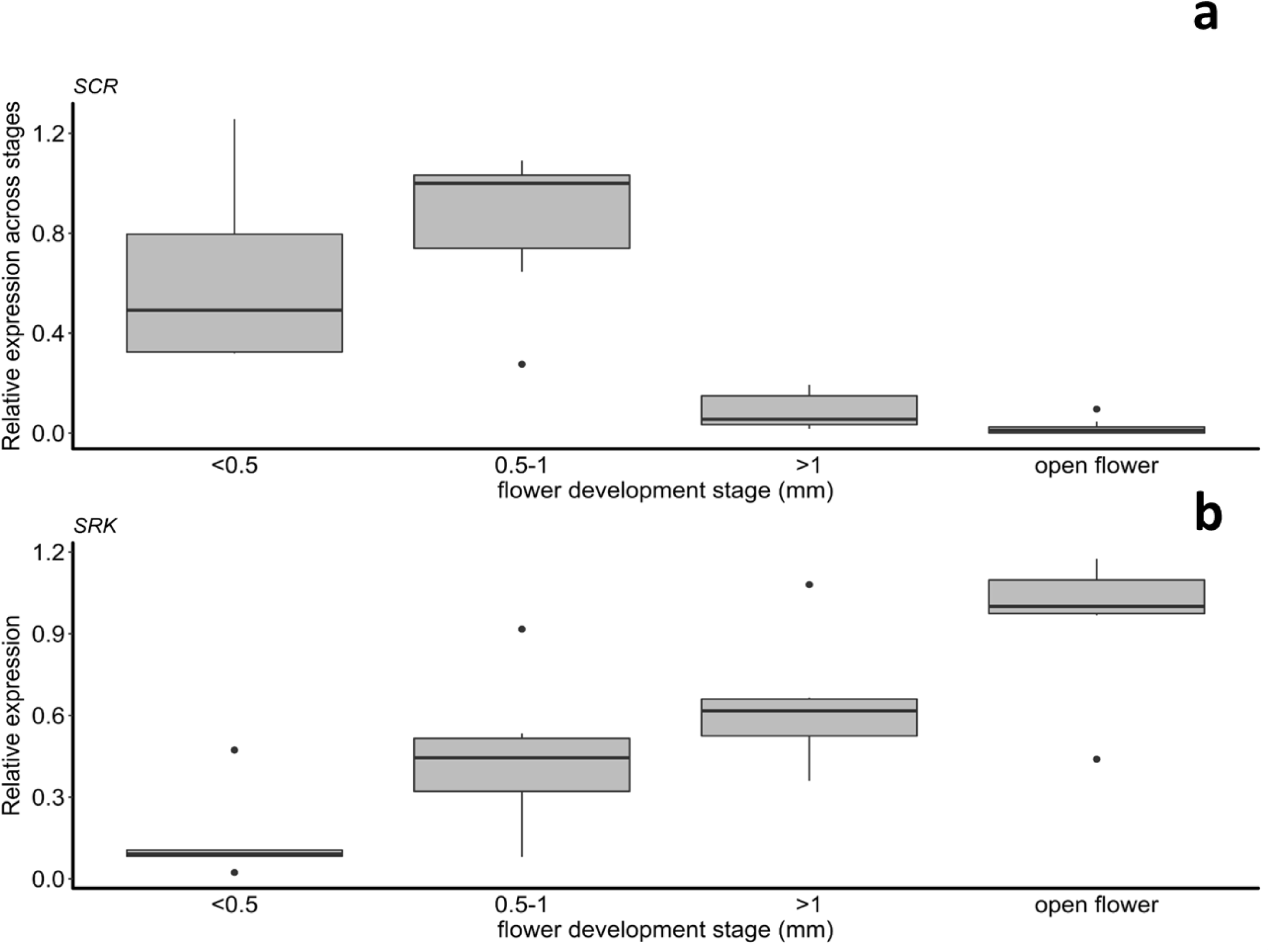
Expression dynamics of **a.** *SCR* and **b.** *SRK* during flower development, from early buds (<0.5mm) to open flowers. For *SCR*, only genotypes in which a given allele was either dominant or co-dominant were included (recessive *SCR* alleles were strongly silenced at all stages and were therefore not informative here). For each allele, 2^−ΔCt^ values were normalized relative to the developmental stage with the highest expression. For each stage, the thick horizontal line represents the median, the box represents the 1^st^ and 3^rd^ quartiles. The upper whisker extends from the hinge to the largest value no further than 1.5 * Inter Quartile Range from the hinge (or distance between the first and third quartiles). The lower whisker extends from the hinge to the smallest value at most 1.5 * IQR of the hinge and the black dots represents outlier values.

### Transcriptional control

Based on these results, we averaged 2^−Δ*Ct*^ values across <0.5mm to 1mm stages to compare expression of a given focal *SCR* allele between genotypic contexts where it was either dominant or recessive relative to the other allele present in the diploid genotype. Of the 54 pairwise interactions for which the dominance phenotype had been firmly established by controlled crosses and the qPCR assay had been performed for both *SCR* alleles (Table S3), as many as 51 (94.4%) are associated with strong asymmetries in transcript levels, with high expression of the dominant *SCR* allele and low expression of the recessive *SCR* allele (Figure 2). Hence, our expression data were largely consistent with the hypothesis of transcriptional control of the dominance hierarchy in pollen genotypic combinations. *SCR* transcripts of the most recessive allele (S1) were only detected in an S1S1 homozygote genotype, but not in any other genotypic combination. Climbing up the dominance hierarchy from most recessive to most dominant, expression of *SCR* was detected in an increasing number of heterozygous combinations, in strong agreement with phenotypic dominance (Figure 2). At the top of the dominance hierarchy, the two most dominant alleles, *SCR*13 and *SCR*20, were expressed in all heterozygous contexts, including when they formed a heterozygote combination with one another (S13S20), also as expected given the codominance observed between them at the phenotypic level (Durand *et al*., 2014). This general rule had a few exceptions however (indicated by arrows on Figure 2). Specifically, we observed low expression for both *SCR*01 and *SCR*12 when in heterozygote combination (S01S12 genotypes) and for both *SCR*10 and *SCR*12 in heterozygote combination (S10S12 genotype), which is not consistent with the documented phenotypic dominance of these alleles in pollen (*S*12> *S*01 and *S*12>*S*10; see Table S3). We also detected expression of both *SCR*02 and *SCR*29 when placed in heterozygote combination, which might explain the unusual phenotypic data indicating robust rejection of pollen from this heterozygote genotype on the [S02] tester line, but only partial compatibility on the [S29] tester line (Table S3). Hence, the dominance interaction between these two alleles may be partial, both at the transcriptional and phenotypic levels. Interestingly, these two alleles belong to class III, which in *A. lyrata* tend to show inconsistent (or leaky) SI responses (Kusaba *et al*. 2001).

**Figure 2:**
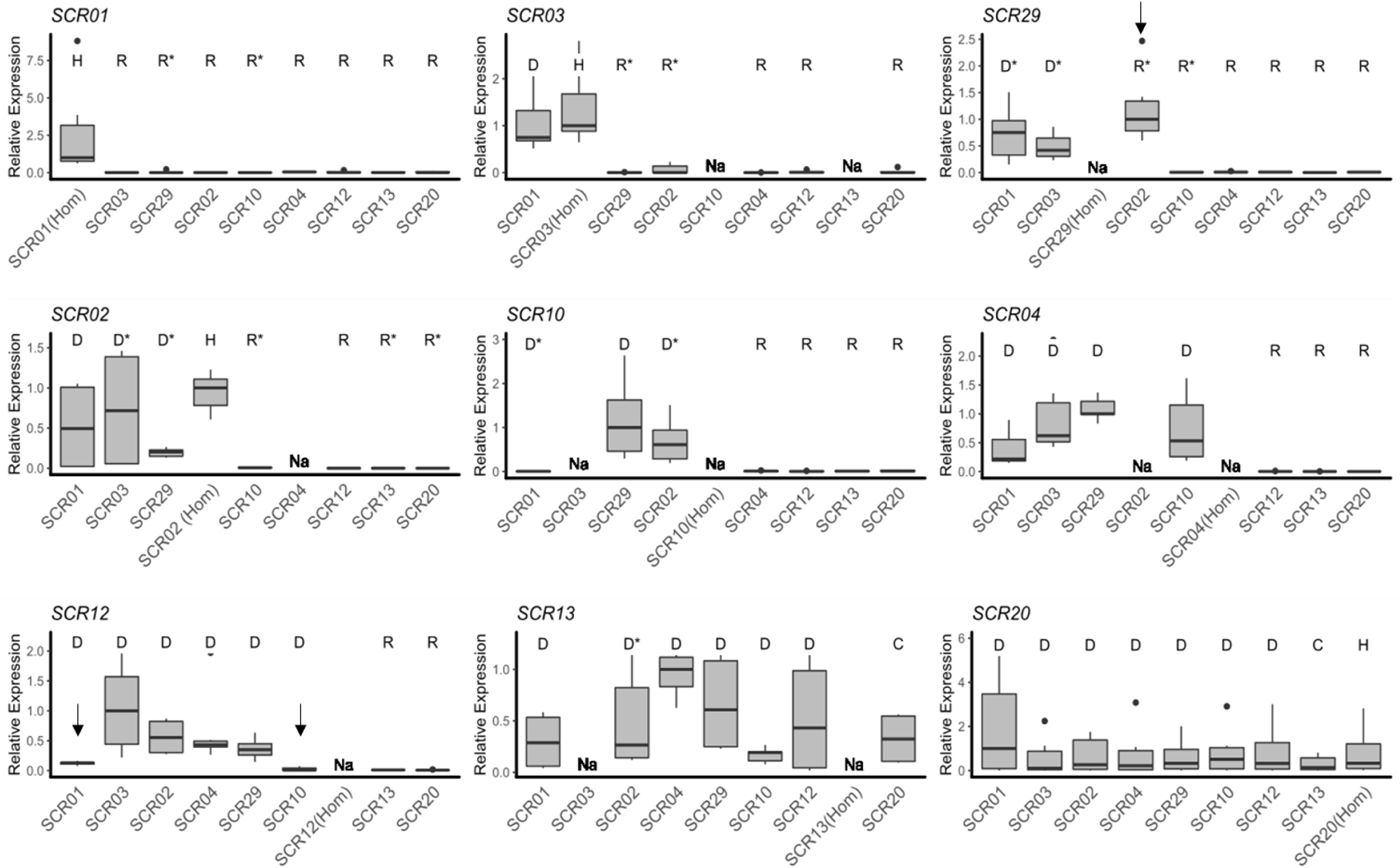
Expression of individual *SCR* alleles in different genotypic contexts. Pollen dominance status of the S-allele whose expression is measured relative to the other allele in the genotype as determined by controlled crosses are represented by different letters (**D**: dominant; **C**: codominant; **R**: recessive; **U**: unknown; **H**: Homozygote, Table S3). In a few instances, relative dominance status of the two alleles had not been resolved phenotypically and were inferred from the phylogeny (marked by asterisks). Thick horizontal bars represent the median of 2^−ΔCt^ values, 1^st^ and 3^rd^ quartile are indicated by the upper and lower limits of the boxes. The upper whisker extends from the hinge to the largest value no further than 1.5 * Inter Quartile Range from the hinge (or distance between the first and third quartiles). The lower whisker extends from the hinge to the smallest value at most 1.5 * IQR of the hinge and the black dots represents outlier values. We normalized values relative to the highest median across heterozygous combinations within each panel. Alleles are ordered from left to right and from top to bottom according to their position along the dominance hierarchy, with SCR01 the most recessive and SCR13 and SCR20 the most dominant alleles. Under a model of transcriptional control of dominance, high expression is expected when a given allele is either dominant or co-dominant and low expression when it is recessive. Exceptions to this model are marked by black vertical arrows and discussed in the text. “Na” marks homozygote or heterozygote genotypes that were not available.

Overall, in spite of these three exceptions, we observed a striking contrast in transcript levels for a given allele according to its relative phenotypic dominance status in the genotype (*F*-value = 19.538; *p*-value < 2.2e-16, Table S5c), suggesting complete silencing of recessive alleles. Specifically, we observed an average 145-fold decrease in transcript abundance in genotypes where a given focal allele was phenotypically recessive as compared to genotypes in which the same focal allele was dominant. We note that the silencing was so strong that the *Ct* values associated with recessive *SCR* transcripts were comparable with those of the negative controls (Figure S3) and close to the detection limits of our method as determined by the break of linearity of the dilution experiment, such that the magnitude of the calculated fold-change value is probably under-estimated (Figure S1). In strong contrast, we found no significant effect of dominance in pistils on *SRK* expression (F-value: 6.8884 p-value: 0.068244; Figure 3, Table S5h), confirming the absence of transcriptional control of dominance for *SRK*.

**Figure 3:**
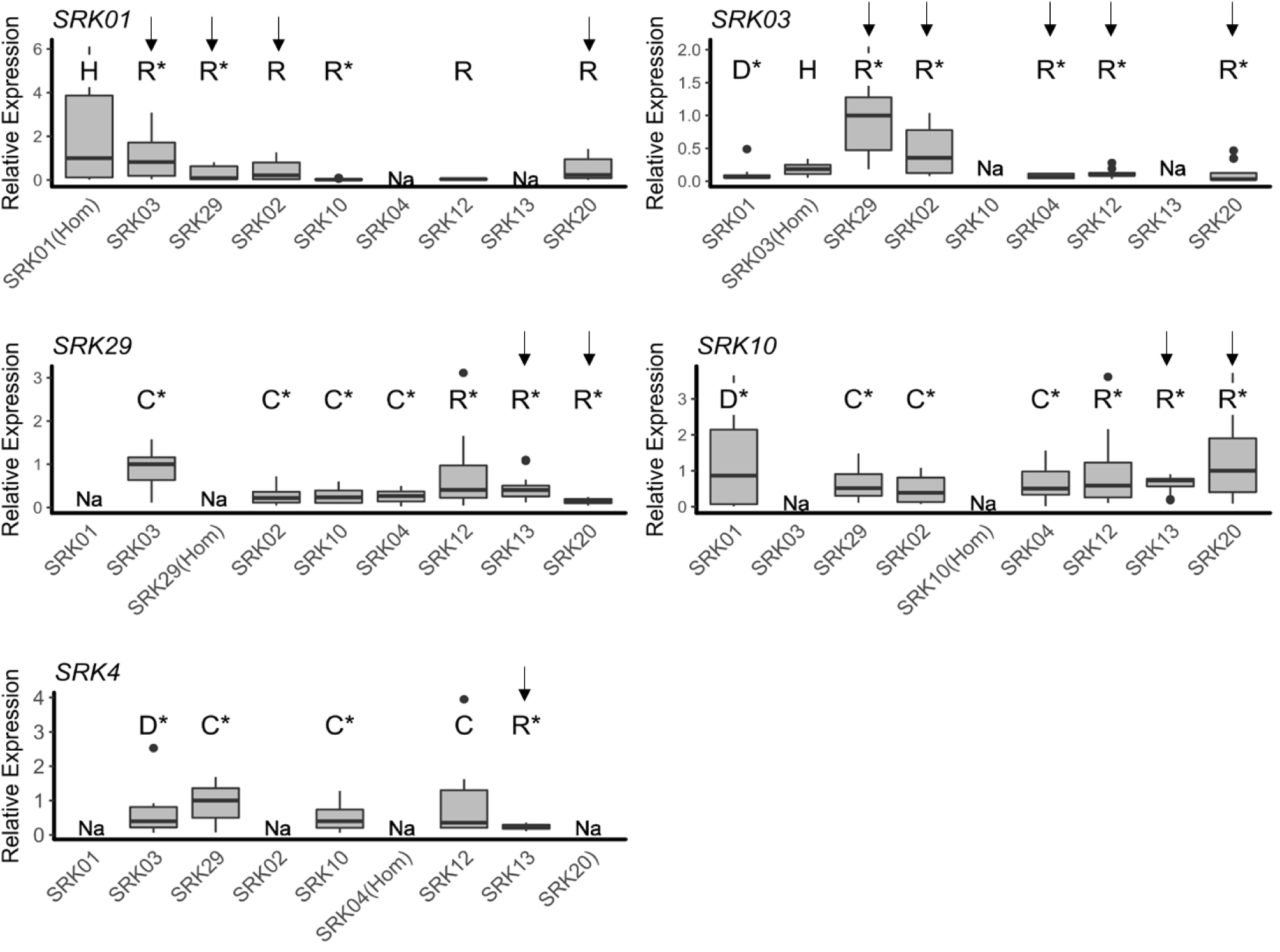
Expression of individual *SRK* alleles in different genotypic contexts. Putative pistil dominance status of the S-allele whose expression is measured relative to the other allele in the genotype is represented by different letters (**D**: dominant; **R**: recessive; **U**: unknown; **H**: Homozygote). Note that the pistil dominance hierarchy of the S-allele have been less precisely determined than the pollen hierarchy, and so many of the pairwise dominance interactions were indirectly inferred from the phylogenetic relationships (and marked by an asterisk) rather than directly measured phenotypically. Thick horizontal bars represent the median of 2^−ΔCt^ values, 1^st^ and 3^rd^ quartile are indicated by the upper and lower limits of the boxes. The upper whisker extends from the hinge to the largest value no further than 1.5 * Inter Quartile Range from the hinge (or distance between the first and third quartiles). The lower whisker extends from the hinge to the smallest value at most 1.5 * IQR of the hinge and the black dots represents outlier values.. We normalized the values for each allele relative to the higher median across heterozygous combination. We normalized values relative to the highest median across heterozygous combinations within each panel. Alleles are ordered from left to right and from top to bottom according to their position in the pistil dominance hierarchy, with SRK01 the most recessive and SRK04 the most dominant allele in our sample, based on the phenotypic determination in Llaurens *et al*. (2008).

### Target features and silencing effect

Levels of *SCR* expression of any given focal allele varied sharply with the alignment score of the “best” target available for the repertoire of canonical sRNAs produced by the other allele present in the genotype (Figure 4a). Specifically, we observed on average high levels of *SCR* transcripts when the score of their best predicted target was low, but consistently low levels of *SCR* transcripts when the score of the best target was high (Figure 4a, Table S5d). Strikingly, the transition between high expression and low expression was abrupt (around an alignment score of 18), suggesting a sharp threshold effect rather than a quantitative model for transcriptional silencing.

**Figure 4:**
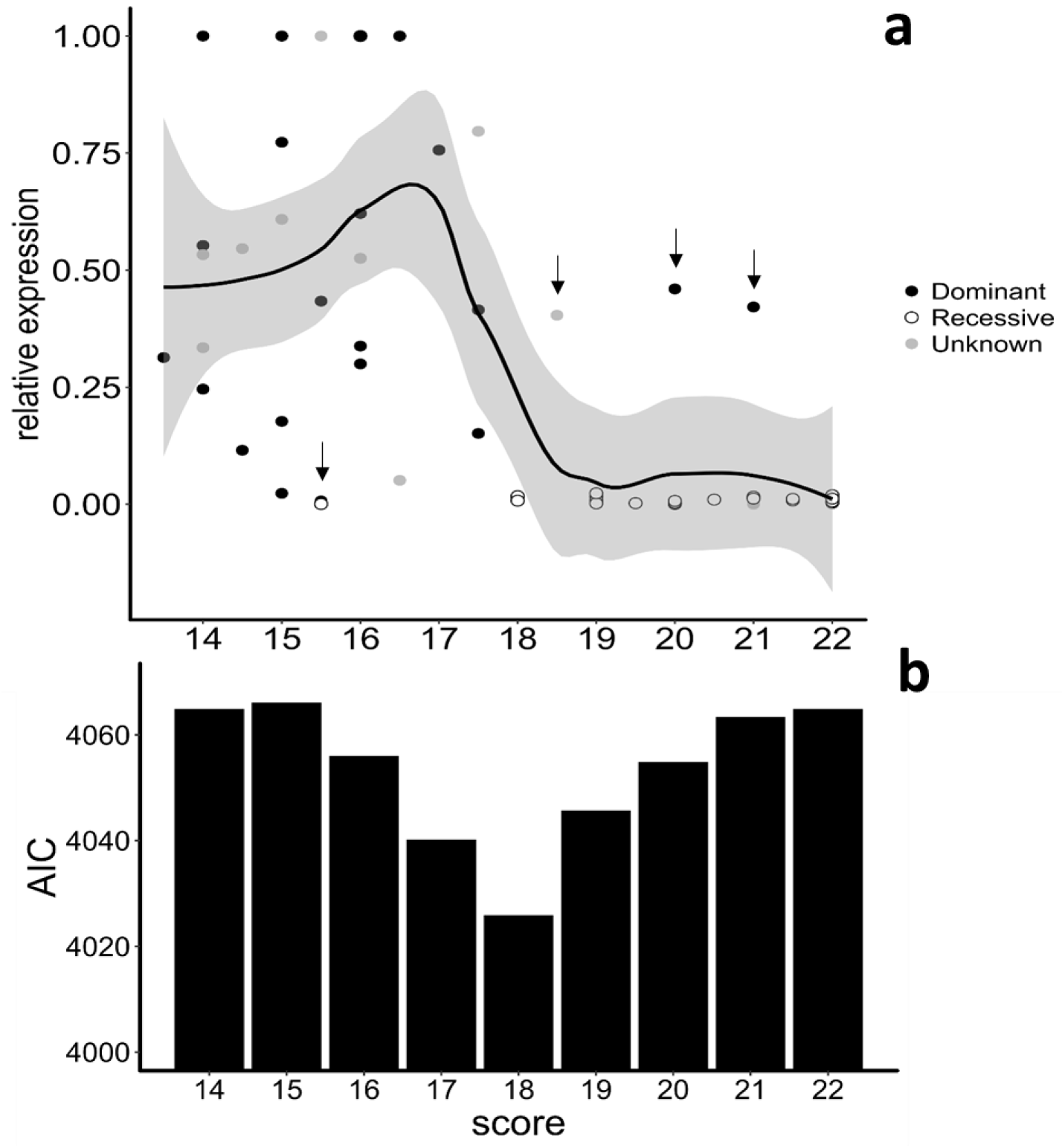
Base-pairing requirements for the transcriptional control of *SCR* alleles by sRNAs suggest a threshold model. **a.** Relative expression of *SCR* alleles as a function of the alignment score of the “best” interaction between the focal allele (including 2kb of sequence upstream and downstream of *SCR*) and the population of sRNAs produced by sRNA precursors of the other allele in the genotype. For each allele, expression was normalized relative to the genotype in which the 2^−ΔCt^ value was highest. Dots are coloured according to the dominance status of the focal *SCR* allele in each genotypic context (black: dominant; white: recessive; grey: undetermined). The black line corresponds to a local regression obtained by a smooth function (loess function, span=0.5) in the ggplot2 package (Wickham, 2009) and the grey area covers the 95% confidence interval. Vertical arrows point to observations that do not fit the threshold model of transcriptional control and are represented individually on Figure 5. **b.** Barplots of the Akaike Information Criteria (AIC) quantifying the fit of the generalized linear model for different target alignment scores used to define functional targets. Lower AIC values indicate a better fit.

In two cases, the presence of a target with a high score within the *SCR* gene of the dominant allele was associated with high relative *SCR* expression (in agreement with the dominant phenotype established by controlled crosses), confirming the absence of silencing (target of Ah04mir4239 on *SCR20*, score=20; and target of Ah10mir4239 on *SCR20*, score =21; Figure 5a) and suggesting that these interactions are not functional. Examining in detail these two exceptions did not reveal mismatches at the 10-11^th^ nucleotide position, suggesting that mismatches at other positions have rendered these sRNA-target interactions inactive (Figure 5a). We note that the target of Ah10mir4239 with the highest score is predicted for a sRNA with non-canonical size (25nt), but this precursor also produces a canonical 24nt isomir with a score above the threshold (score = 20, Table S6). These two sRNAs (Ah04mir4239 and Ah10mir4239) have a 5’ nucleotide different from the expected “A” for 24nt sRNAs, possibly suggesting that loading into an improper AGO protein may have rendered these predicted interactions inactive. Another exception concerns the observed low score (15.5) for the best match between a sRNA from the dominant allele Ah04mirS4 and its best putative target at the recessive *SCR03* (Figure 5b). Whether *SCR*04 silences *SCR03* through this unusual target or through another elusive mechanism remains to be discovered.

**Figure 5:**
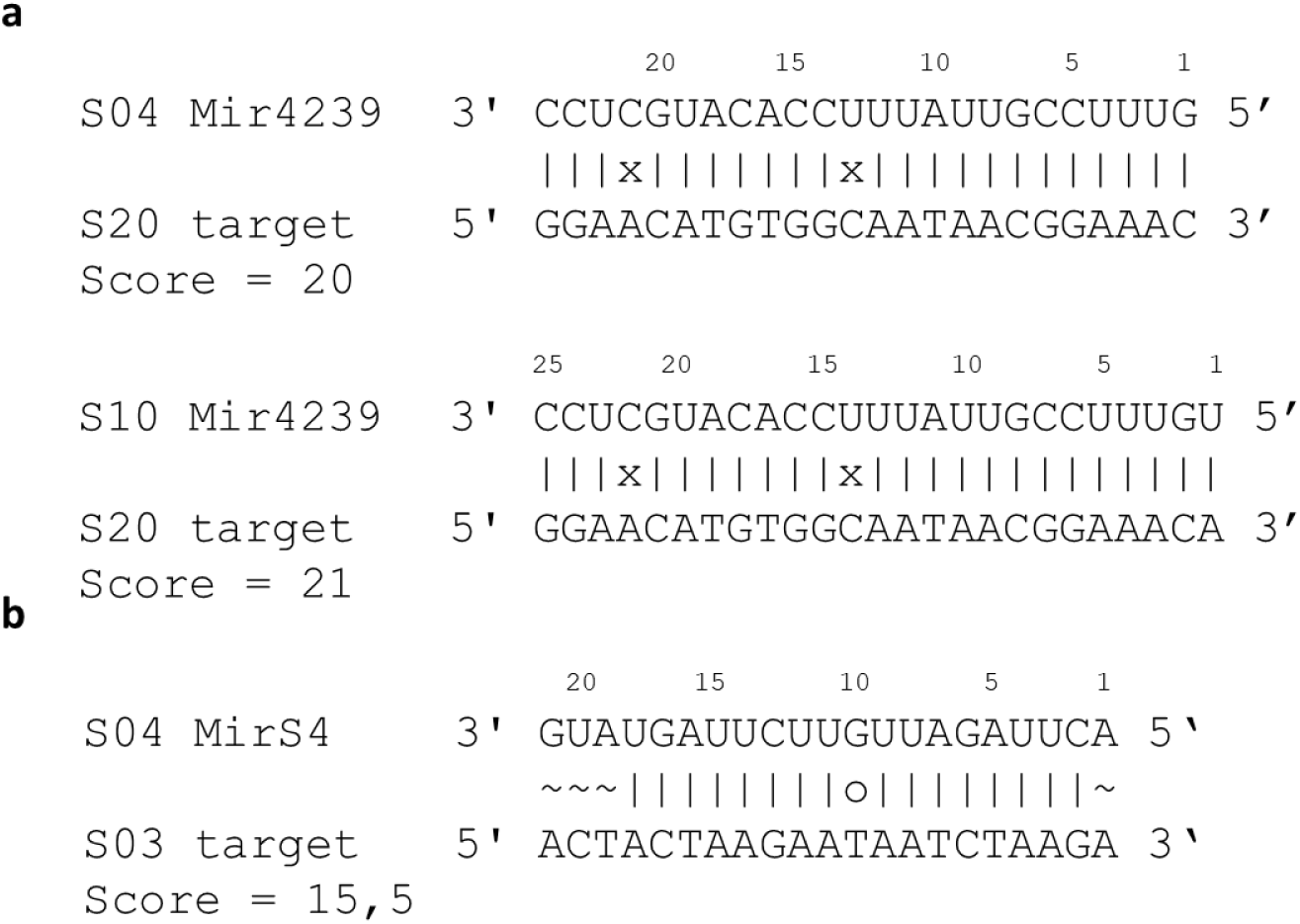
Predicted sRNA/target interactions that do not fit with the documented dominance phenotype or the measured expression. For each alignment, the sequence on top is the sRNA and the bottom sequence is the best predicted target site on the *SCR* gene sequence (including 2kb of sequence upstream and downstream of *SCR*). **a.** sRNA targets with a score above 18, while the S-allele producing the sRNA is phenotypically recessive over the S-allele containing the *SCR* sequence. **b.** sRNA target with a score below 18, while the S-allele producing the sRNA (S04) is phenotypically dominant over the S-allele containing the *SCR* sequence and transcript levels of the *SCR03* allele is accordingly very low. This is the best target we could identify on SCR03 for sRNAs produced by S04.

In spite of the generally very low expression of all recessive alleles, we found marginal evidence that the strength of silencing experienced by a given *SCR* allele varies across genotypic combinations for a given allele (F-value=2.221, *p*-value = 0.0756, Table S5i). However, there was no evidence that the position of the target site on the measured allele (promoter; intron; intron-exon boundary; upstream *vs.* downstream) could explain this variation (F-value=1.7061, *p*-value = 0.1928, TableS5e). We also found no effect of the inferred age of the miRNA on the mean alignment score (mean= 20.41 and 20.22 for recent or ancient miRNAs, respectively; F-value: 0.0362; *p*-value = 0.8504, Table S5j). Finally, we compared the alignment scores observed here in Arabidopsis with those in Brassica for *Smi* & *Smi2* on their *SCR* target sequences. A clear threshold was also observed, but in Brassica the alignment score threshold distinguishing dominant from recessive interactions was 16.5 instead of 18 (Table S6), suggesting distinct base-pairing requirements for effective silencing in these two systems.

## Discussion

Determining the base-pairing requirement for sRNA silencing in plants has remained challenging because the “rules” used for target prediction have typically been deduced from observations that conflate distinct microRNA genes and their distinct mRNA targets over different genes. Moreover, detailed evaluations of the functional consequences of mismatches have relied on heterologous reporter systems (typically GFP in transient tobacco assays), hence limiting the scope of the phenotypic consequences that can be studied. Here, we build upon the inter-allelic regulatory system controlling transcriptional activity of alleles of the SI system in Arabidopsis revealed in Durand *et al*. (2014), where multiple sRNAs regulate target sites on alleles of a single gene (*SCR*), and in which we are able to make a direct link between the sRNA-target interactions, the level of *SCR* transcript and the encoded phenotype (dominance/recessivity interaction). The first step was to clarify several aspects of the expression pattern of the genes controlling SI in *A. halleri*, which was necessary to confirm that the dominance interactions in Arabidopsis involve transcriptional regulation.

### Expression profile

Earlier accounts had suggested that alleles of the allelic series may differ from one another in their expression profile (Kusaba *et al*., 2002). In line with Kakizaki *et al*., (2003), Suzuki *et al.*, (1999); Schopfer *et al*., (1999); Takayama *et al*., (2000) and Shiba *et al*., (2002), we found maximal expression of *SCR* in early buds but low or no expression at the open flower stage. This expression pattern is consistent with in situ hybridization experiments showing that SCR transcripts are localized in the tapetum, a specialized layer of cells involved in pollen grains coating (Iwano *et al*., 2003) which undergoes apoptosis and is quickly degraded as the development of pollen grains inside the anther progresses (Murphy & Ross, 1998; Takayama *et al*., 2000). We confirmed that differences exist in the temporal dynamics of expression among alleles, as suggested by Kusaba *et al*. (2002) in *A. lyrata*, possibly as the result of strong sequence divergence of the promotor sequences of the different *SCR* alleles. Finally, we confirmed that *SCR* and *SRK* have sharply distinct expression dynamics throughout flower development. Indeed, transcript levels of *SRK* increased steadily along development and were very low in early buds, consistent with the observation that SI can be experimentally overcome to obtain selfed progenies by “bud-pollination” (Llaurens *et al*. 2009).

### Generality of the transcriptional control of dominance in Arabidopsis

Based on this clarified transcriptional dynamics, we confirmed the generality of the transcriptional control of dominance for *SCR*, with as much as 96.3% of the documented dominance interactions associated with mono-allelic expression of the dominant *SCR* allele and complete silencing of the recessive *SCR* allele in heterozygote genotypes. Even in the single heterozygote genotype where in our previous study (Durand *et al*., 2014) no sRNA produced by the phenotypically dominant allele was predicted to target the sequence of the phenotypically recessive *SCR* allele (e.g. S04>S03), transcripts from the recessive *SCR03* allele were undetected. This suggests either that some functional sRNAs or targets have remained undetected by previous sequencing and/or by our *in silico* prediction procedures, or that mechanisms other than sRNAs may cause transcriptional silencing for some S-allele combinations. Regardless of the underlying cause, the generality of the transcriptional control of dominance suggests that the simple comparison of transcript levels between the two alleles in a heterozygote genotype could be used as a first approximation to determine their relative dominance levels. In contrast, we confirmed the absence of transcriptional control for *SRK*, for which both alleles were consistently expressed at similar levels in all heterozygote genotypes examined, irrespective of the (pistil) dominance phenotype. For *SRK*, other dominance mechanisms must therefore be acting, which are yet to be discovered (*e.g.* Naithani *et al*., 2007).

### Variation in the strength of silencing

An important feature of the silencing phenomenon is that the decrease of transcript levels for recessive *SCR* alleles was very strong in heterozygous genotypes, bringing down transcript levels below the limits of detection in most cases. This is in line with the intensity of transcriptional silencing by heterochromatic siRNAs (typically very strong for transposable element sequences, see Marí-Ordóñez *et al*., 2013), while post-transcriptional gene silencing by microRNAs can be more quantitative (Liu *et al*., 2014). As a result of this strong decrease of transcript levels, the strength of silencing appeared independent from the position of the sRNA target along the *SCR* gene (promoter *vs.* intron), although we note that our power to distinguish among levels of transcripts of recessive alleles, which were all extremely low, is itself fairly low. It remains to be discovered whether the different positions of the sRNA targets (Durand *et al*., 2014) do indeed imply different transcriptional silencing mechanisms.

### A simple threshold model for sRNA-based silencing

Based on the many allelic combinations where we could compare the agnostic prediction of putative target sites with the level of transcriptional silencing, we find that a simple threshold model for base-pairing between sRNAs and their target sites captures most of the variation in *SCR* expression in heterozygotes. This result provides a direct experimental validation of the *ad-hoc* criteria used in Durand *et al*., (2014). However, our results also indicate that this quantitative threshold is not entirely sufficient to capture the complexity of targeting interactions. Indeed, in three of the 54 cases tested this simple threshold model would inappropriately predict targeting of a dominant *SCR* allele by a sRNAs from a more recessive allele, yet the dominant *SCR* allele was expressed at normal levels with no sign of silencing in these heterozygote genotypes (Figure 5a). The targeting interaction may be abolished either by defects in the sRNA itself (e.g. for Ah04mir4239 the 5’ nucleotide is a G, while the majority of functional 24 nt small RNA molecules end with a 5’A, which may interfere with loading in the appropriate AGO protein). Alternatively, the targeting interactions may be abolished by the position of the mismatches (at position 14 and 22 of the Ah10mir4239 and at position 13 and 21 of the Ah04mir4239, both on *SCR20*).

Similarly, a single mismatch at position 10 in the *Smi* interaction in Brassica (Tarutani *et al*., 2010) and in other microRNA-targets interactions (Franco-Zorrilla *et al*., 2007) was shown to result in loss of function of the interaction (Table S6). Interestingly, quantitative differences may exist between Arabidopsis and Brassica, as the experimentally validated targets in Brassica (Tarutani *et al*., 2010; Yasuda *et al*., 2016) correspond to base-pairing threshold below the one that we find in Arabidopsis (*i.e.* a target score of 16.5 seems sufficient for silencing in Brassica *vs.* 18 in Arabidopsis). For Brassica, both class I and class II alleles have *Smi*, but a mismatch at the 10^th^ position was proposed to explain why the class II *Smi* is not functional. Here, we show that this mismatch drives the alignment score below the 16.5 threshold and could be sufficient to explain the loss of function, regardless of its position. Overall, although these small RNAs achieve their function in a way that may be sharply different from classical microRNAs (DNA methylation *vs.* mRNA cleavage), our results suggest that the sRNA-target complementarity rules for silencing in both cases are qualitatively consistent (Liu *et al*., 2014). Better understanding the molecular pathway by which these sRNAs epigenetically silence their target gene (*SCR*) will now be key to determine whether this threshold model can be generalized to more classical siRNAs found across the genome, as evidence is still missing for such classes of sRNAs.

### Implications for the evolution of the dominance hierarchy

The existence of a threshold model has important implications for how the dominance hierarchy can evolve. In fact, our model suggests that a single SNP can be sufficient to turn a codominance interaction into a dominance interaction (and vice-versa), making this a relatively trivial molecular event. This is actually what Yasuda *et al*., (2016) observed in *B. rapa*, where the combination of single SNPs at the sRNA *Smi2* and its *SCR* target sequences resulted in a linear dominance hierarchy among the four class II S-alleles found in that species. Strikingly, in some cases, we observed base pairing at sRNA-target interactions with very high alignment scores (up to 22), *i.e.* above the threshold at which transcriptional silencing was already complete (score =18). Under our simple threshold model, such interactions are not expected since complete silencing is already achieved at the threshold, and no further fitness gain is therefore to be expected by acquiring a more perfect target. A first possibility is that these interactions reflect the recent emergence of these silencing interactions. In fact, one of the models for the emergence of new microRNAs in plant genomes involves a partial duplication of the target gene, hence entailing perfect complementarity at the time of origin that becomes degraded over time by the accumulation of mutations (Allen *et al*., 2004). Under this scenario, the higher-than-expected levels of sRNA-target complementarity could reflect the recent origin of these sRNAs but we found no evidence of a difference in alignment score for young *vs.* old sRNA precursors. A second possibility is that selection for developmental robustness is acting to prevent the phenotypic switch from mono- to bi-allelic expression of *SCR* (especially during stress events, Boukhibar & Barkoulas, 2016) that could be devastating for the plant reproductive fitness (Llaurens *et al*. 2009). Indeed, we observed strong variation in overall *SCR* expression when the sRNA target score of the companion allele is below the threshold in the benign greenhouse conditions under which we grew our plants, and it is possible that under stress conditions the epigenetic machinery may be less efficient, hence requiring stronger base-pairing to achieve proper silencing. Finally, a third possibility is that sRNA-target complementarity above the threshold reflects the pleiotropic constraint of having a given sRNA from a dominant allele control silencing of the complete set of target sequences from the multiple recessive alleles segregating, and reciprocally of having a given *SCR* target in a recessive allele maintaining molecular match with a given sRNA distributed among a variety of dominant alleles. Comparing the complementarity score of sRNA/target interactions among sRNAs or targets that contribute to high versus low numbers of dominance/recessive interactions will now require a more complete depiction of the sRNA-target regulatory network among the larger set of S-alleles segregating in natural populations.

## Acknowledgments

We thank Sylvain Billiard and Isabelle de Cauwer for statistical advice and discussions, Romuald Rouger and Anne Duputié for help with producing figures and Alexis Sarazin and three anonymous reviewers for comments on the manuscript. This work was funded by the European Research Council (NOVEL project, grant #648321). N.B. was supported by a doctoral grant from the president of Université de Lille-Sciences et Technologies and the French ministry of research. The authors also thank the Région Hauts-de-France, and the Ministère de l’Enseignement Supérieur et de la Recherche (CPER Climibio), and the European Fund for Regional Economic Development for their financial support

## Author Contribution

NB, SS, SB, ACH performed the molecular biology experiments. CP and ES obtained and took care of the plants. SS, IFL and XV provided advice on the experimental strategy and interpretations. NB performed the statistical analyses. VC supervised the work. NB and VC wrote the manuscript.

---

**Figure S1.**
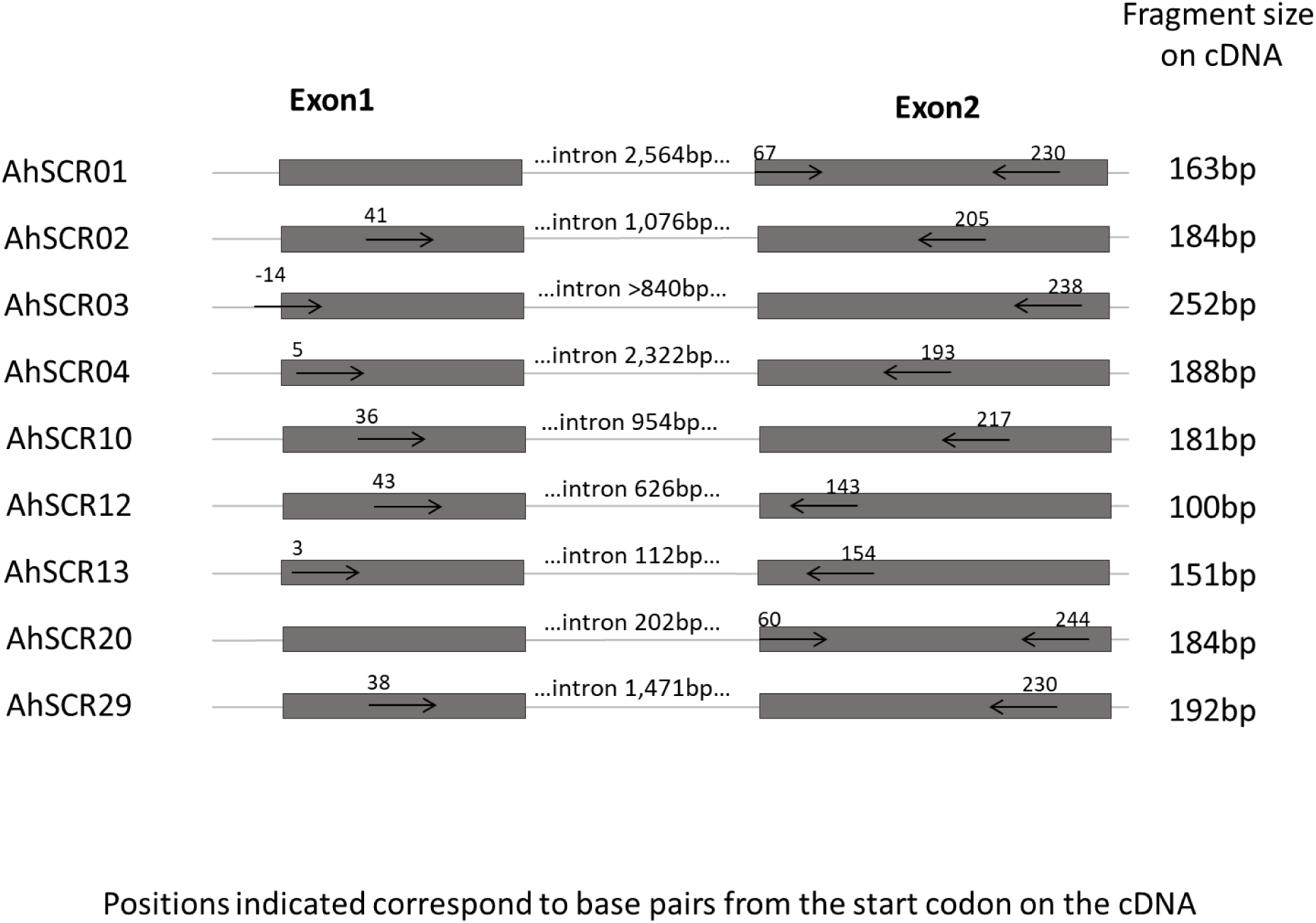
Position of the *SCR* qPCR primers. Each primer is represented by a black arrow, according to its relative position to the gene. Positions indicated correspond to base pairs from the start codon on the cDNA.

**Figure S2.**
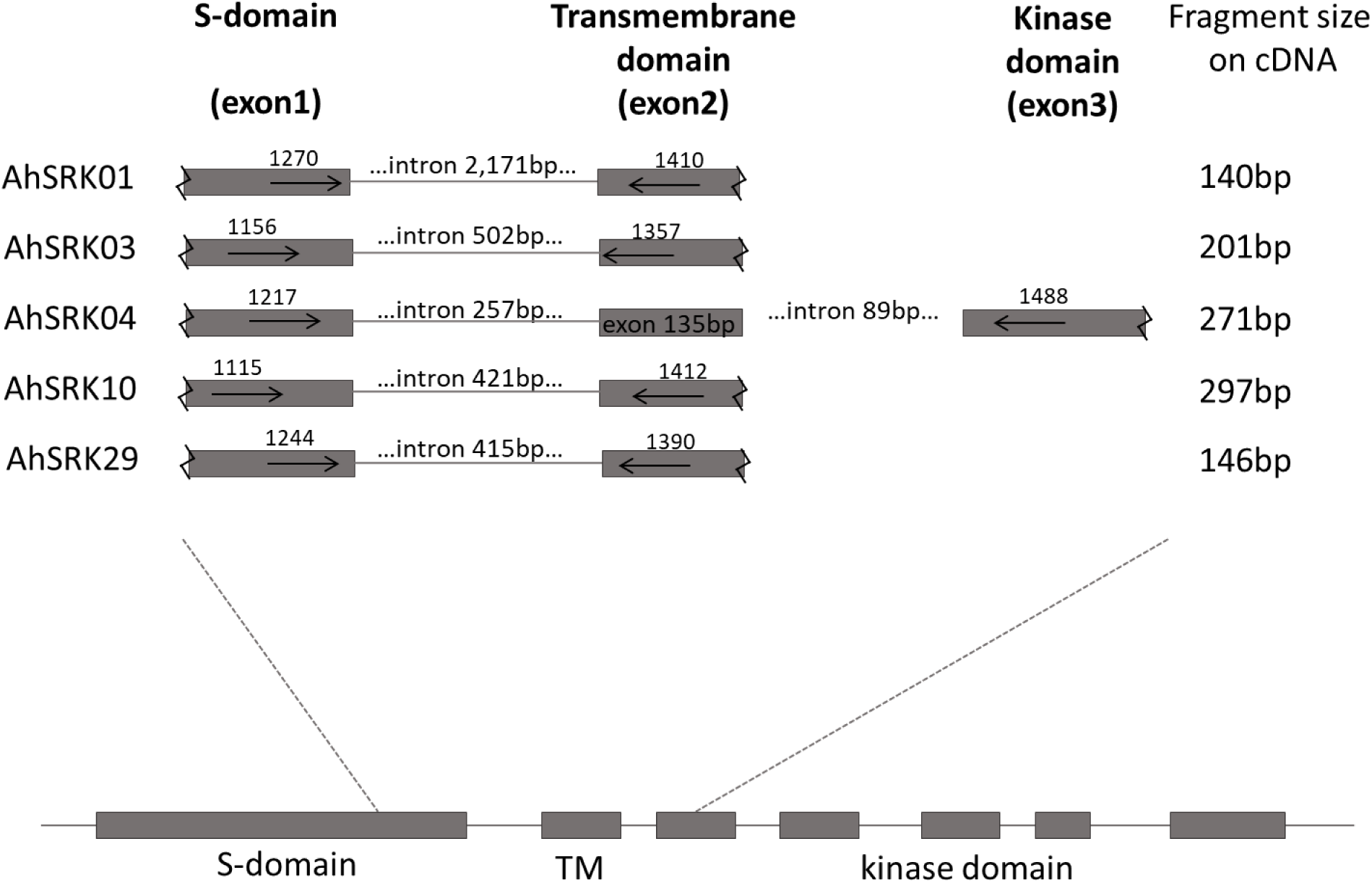
Position of the *SRK* qPCR primers. Each primer is represented by a black arrow, according to its relative position to the gene. Positions indicated correspond to base pairs from the start codon on the cDNA.

**Figure S3.**
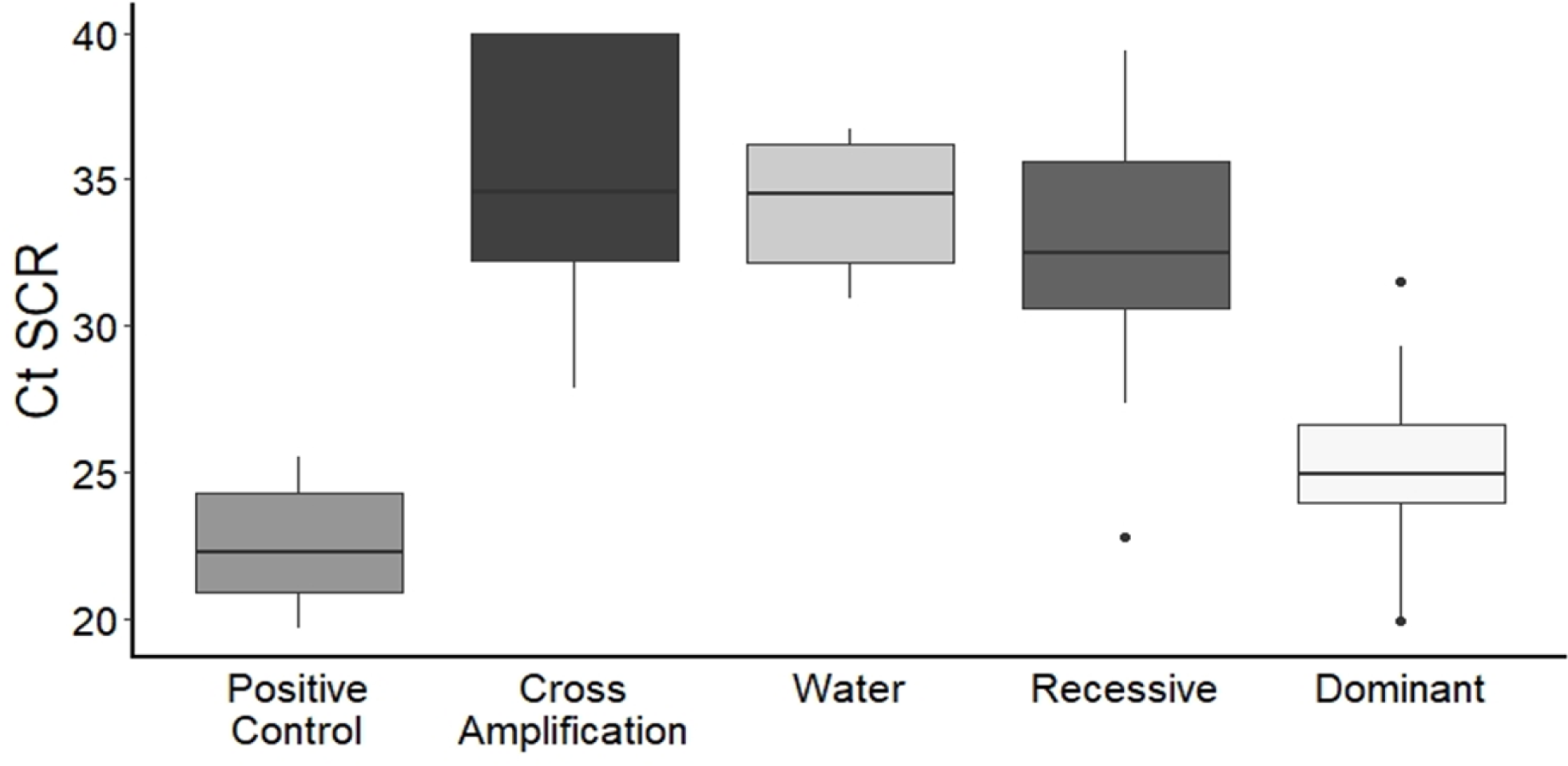
Validation of the *SCR* qPCR primers. “Positive control**”**: amplification assay with primers for *SCR* alleles that are expressed in the cDNA used. “Cross Amplification”: amplification assay with primers for *SCR* alleles that are different from the ones in the cDNA used. “Water**”** : amplification assay with primers but water instead of cDNA. “Recessive”: amplification with primers for the phenotypically recessive *SCR* allele in the cDNA (mean *Ct* values of the biological and technical replicates). “Dominant”: amplification with primers for the phenotypically dominant *SCR* allele in the cDNA (mean *Ct* values of the biological and technical replicates). Thick horizontal bars represent the median of 2^−ΔCt^ values, 1^st^ and 3^rd^ quartile are indicated by the upper and lower limits of the. The upper whisker extends from the hinge to the largest value no further than 1.5 * Inter Quartile Range from the hinge (or distance between the first and third quartiles). The lower whisker extends from the hinge to the smallest value at most 1.5 * IQR of the hinge and the black dots represents outlier values.

**Figure S4:**
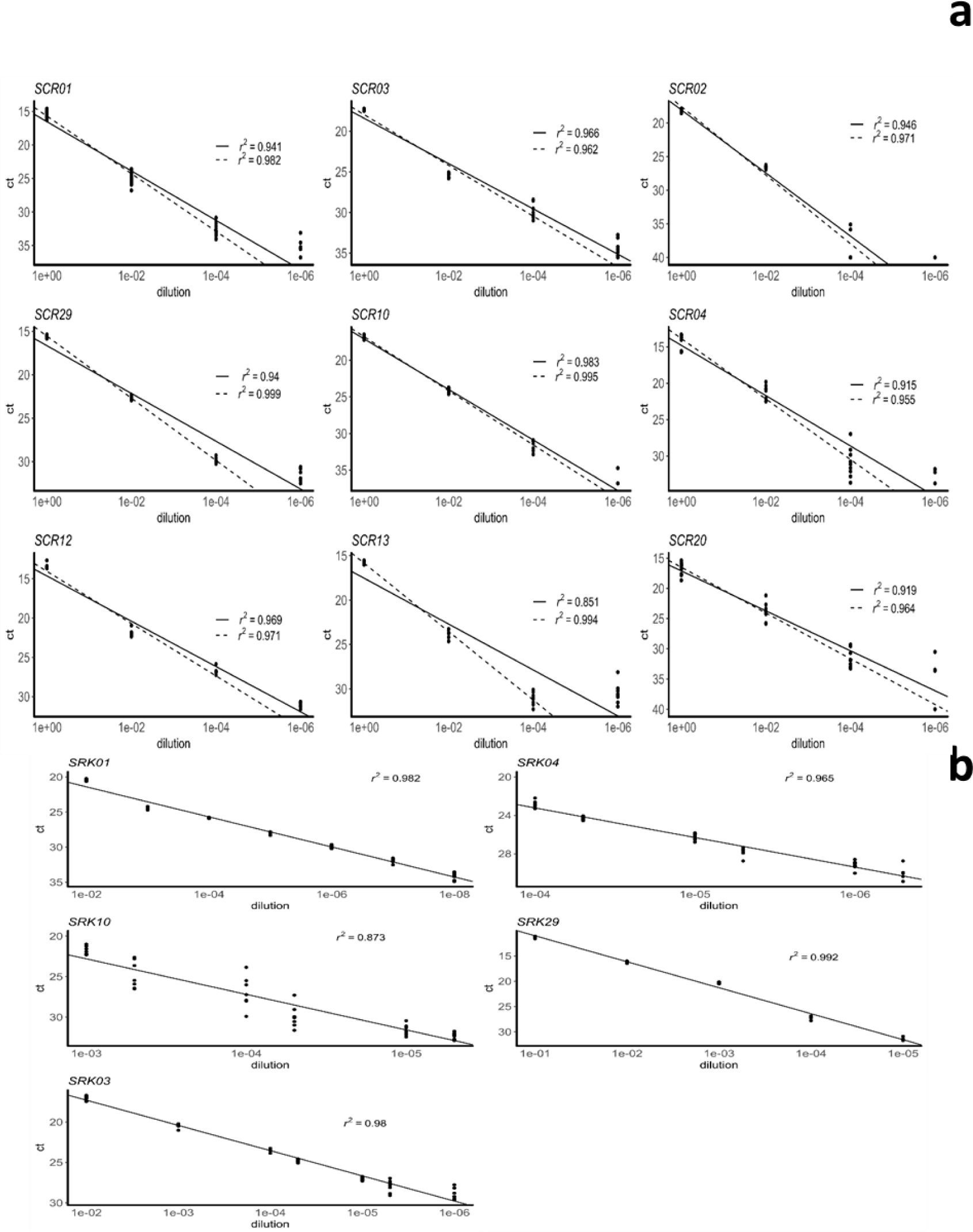
qPCR amplification (non-transformed *Ct* values) in serial dilutions for each *SCR* (**a**) and *SRK* (**b**) allele. Solid lines are the linear regressions over all *Ct* values. Dashed lines are linear regressions excluding the highest dilution level.

**Figure S5.**
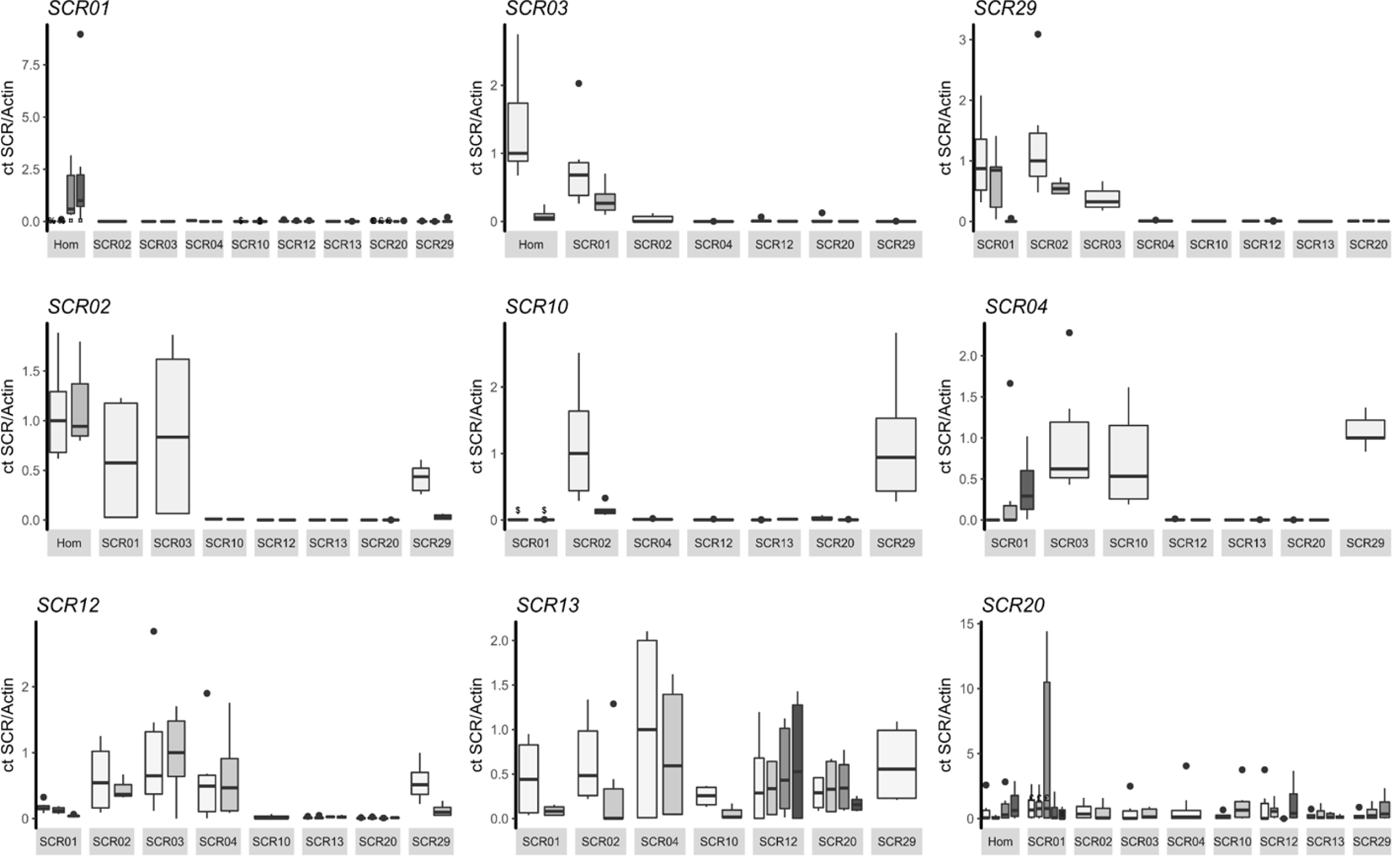
Expression of individual *SCR* alleles in different genotypic contexts, representing each biological and clone replicate separately. Symbols on top of the boxes indicate measures from identical clone replicates. See legend of Figure 2 for a full description.

**Figure S6.**
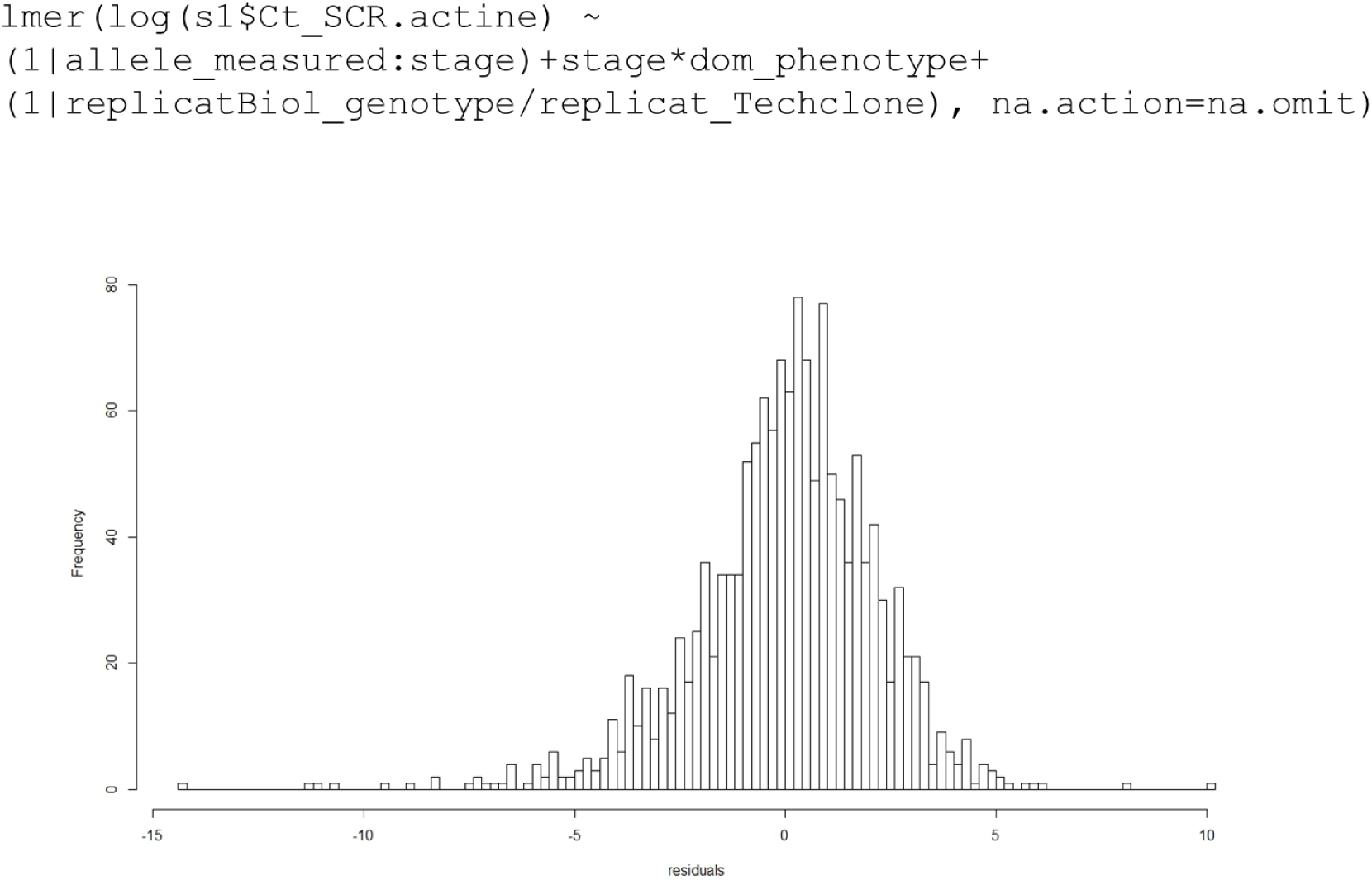
Generalized linear mixed model used to test the effect of developmental stage and dominance status on the expression of *SCR* alleles (*Ct* values). The distribution shows that the residues of the full model are approximately normally distributed when taking allele identity, developmental stage and dominance status into account and using a logarithmic transformation of the *Ct*_SCR_ /*Ct*_actin_ ratios.

**Table S1**. *SCR* samples analysed for each S-locus genotype, showing the number of biological and clone replicates over the four developmental stages sampled. “Allele 1” refers to the first allele noted in the genotype (for example in the S1S2 genotype, “allele 1” is S1 and “allele 2” is S2).

**Table S2**: *SRK* samples analysed for each S-locus genotype, showing the number of biological and clone replicates over the four developmental stages sampled. The alleles are named accordingly to the Table S1.

**Table S3**: Dominance relationships between alleles from the different genotypes included in this study as determined by controlled crosses.

**Table S4**: qPCR primer sequences for each *SCR* and *SRK* alleles studied.

**Table S5**: Detailed results from the generalized linear mixed models. **a**. Decomposition of the sources of variance across allele identity and the hierarchical levels biological, clones and technical replicates for *SCR*. **b**. Test of the variation of expression dynamic across *SCR* alleles. **c**. Test of the dominance and stage effects on *SCR* transcript levels, showing a significant interaction. **d**. Comparison of the fit of the model under different base-pairing score thresholds. **e**. Test of the effect of the position of the target on the strength of silencing. **f**. Decomposition of the source of variance across the technical replicates and the allele identity for *SRK*. **g**. Test of the variation of expression dynamic across *SRK* alleles. **h.** Test of the effect of stage and dominance on *SRK* transcript levels. **i**. Test of the effect of the identity of the companion allele on *SCR* transcript levels. **j**: Test of the effect of age on alignment score above the threshold of 18.

**Table S6**: sRNA and *SCR* target identified as the best match for every pair of S-alleles. *Ct_SCR_/Ct_actin_* ratios are given for the target S-allele in the interaction and is calculated as the mean value across the two earliest developmental stages (see Figure 1). The positions of the targets are given relative to the beginning of the nearest exon of *SCR* for targets upstream from the gene or in the intron, and relative to the stop codon for downstream targets. R: Recessive; D: dominant as phenotypically determined by controlled crosses; H: homozygote; R* or D*: dominance as indirectly inferred from the phylogeny of *S*-alleles. The size of the canonical (21 or 24nt-long) isomiR with the highest targeting score is given in parentheses. The targeting score of the best canonical isomiR is given in parentheses).

